# Finite Element Modeling of Electrical Activity in Human Uterine Tissue: Advances in Simulation Techniques

**DOI:** 10.1101/2024.10.06.616865

**Authors:** Saeed Zahran

## Abstract

Simulating the electrical activity of the human uterus has become a crucial tool for understanding the underlying biophysical phenomena, such as uterine contractions during pregnancy. Finite element models (FEM) offer valuable insights into these dynamics by providing a scalable framework to explore the propagation of electrical signals at the cellular and tissue levels. This study presents a finite element-based bidomain model to simulate the excitation propagation across the human uterus. Our model integrates cellular-level electrophysiological properties with tissue-level electrical propagation using the FEniCS Python library. A three-dimensional, realistic representation of uterine tissue is employed to simulate excitation patterns, contributing to a deeper understanding of uterine electrophysiology. The model can also be adapted to investigate pathological conditions such as preterm labor and test potential interventions. The developed simulation framework provides a scalable solution for the numerical challenges posed by solving complex, non-linear ordinary differential equations (ODEs) associated with uterine electrical activity. These simulations could offer a foundation for future research on uterine function and its related disorders.

## I. Introduction

The study of electrical activity in the uterus has emerged as a critical area of research, especially in understanding uterine behavior during pregnancy and labor. Computational modeling of this activity provides invaluable insights into how uterine smooth muscle cells behave under various physiological conditions. Uterine contractions, which are vital during labor, are driven by synchronized electrical signals among these muscle cells. Simulating this process computationally offers an efficient means to explore the underlying mechanics, detect abnormalities, and potentially predict or prevent premature births [1].

Finite Element Models (FEM) have become indispensable in biomedical research due to their ability to solve complex, multi-scale problems with high precision [2]. The uterus, like other excitable tissues, presents unique challenges in modeling due to its non-linear biophysical properties and the interaction between intracellular and extracellular domains. Traditional approaches, such as modeling the uterus as a simple geometric structure (e.g., spherical or ellipsoidal), fail to capture the full complexity of its behavior. Recent advancements in imaging technologies, particularly MRI-based meshing, now allow for the creation of highly realistic three-dimensional (3D) uterine models. These models provide a more accurate platform for simulating electrical signal propagation in uterine tissue [3].

The propagation of electrical activity in the uterus can be described using reaction-diffusion equations, specifically the bidomain model. Initially developed for cardiac tissue modeling, the bidomain model has been adapted to represent the uterus [4]. This model divides the tissue into two continuous domains: intracellular and extracellular spaces. The electrical potential differences between these domains drive the excitation of smooth muscle cells, which, when coordinated, lead to uterine contractions. By solving systems of partial differential equations (PDEs), the bidomain model enables the simulation of spatial and temporal propagation of electrical signals throughout the uterus [5].

At the cellular level, these electrical activities are represented by action potentials (AP) driven by ion exchanges across the cell membrane. In this study, we integrate the Red3 electrophysiological model, which was specifically developed to simulate the action potentials of human uterine tissue. The Red3 model captures key ionic currents, including calcium and potassium, which are essential for electrical excitability and propagation of action potentials [6]. This integration of cellular and tissue-level models provides a comprehensive framework for studying both normal and pathological uterine behavior, such as labor or preterm birth.

With the rising prevalence of preterm labor globally, the demand for accurate modeling tools capable of predicting abnormal uterine behavior has grown. Recent studies, such as those by Yochum et al. [1] and La Rosa et al. [8], emphasize the potential of computational models in simulating uterine contractions and their impact on labor. Furthermore, recent works, including our own contributions on sparse source imaging [12], source localization using the maximum entropy on the mean approach [14], and the separation of electrohysterogram (EHG) sources using tensor models [15], have introduced robust models capable of providing more realistic simulations of electrical activity across the human uterus. Our study specifically introduces a 3D finite element framework using the FEniCS Python library to enable scalable and accurate simulations of uterine electrical activity, addressing both normal and pathological conditions.

In this paper, we present a comprehensive finite element model of uterine electrical activity, applying the bidomain framework to simulate excitation across a 3D realistic uterine tissue model. This approach offers an advantage by using realistic uterine geometry, contrasting with earlier models that relied on simplified shapes. By refining the mathematical framework and incorporating recent advancements in FEM solvers, this study aims to provide a flexible and scalable tool for research and potential clinical applications [9].

Our approach not only addresses the complexity of solving non-linear systems of ordinary differential equations (ODEs) and PDEs but also introduces novel techniques to model the anisotropic propagation of electrical signals across distinct regions of the uterus. This allows for detailed analysis of how different regions of the uterus coordinate during labor and how disruptions in this coordination could lead to complications such as preterm birth [11].

The results demonstrate the efficacy of our approach in simulating electrical activity in the uterus. Our model offers a foundation for future research into both physiological and pathological uterine behavior, providing potential applications in personalized medicine. By creating patient-specific uterine models, the method could help predict labor patterns and guide medical interventions in clinical settings [12], [13], [14], [15].

## II. Materials and methods

### A. Action Potential Model

Uterine smooth muscle cells are excitable cells that propagate electrical signals through the tissue via gap junctions, allowing for synchronized contractions [6]. To model this, we used the Red3 model, a detailed electrophysiological model representing ion channel dynamics in uterine muscle cells. The Red3 model accounts for ionic currents, including voltage-dependent calcium (ICa), potassium (IK), calcium-activated potassium (IKCa), and leakage (IL) currents.

The evolution of the transmembrane potential Vm is governed by:

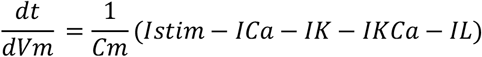

where *ICa, IK, IKCa* and IL represent the individual ionic currents, Cm is the membrane capacitance, and Istim is the external stimulus current. The individual ionic currents are modeled as:

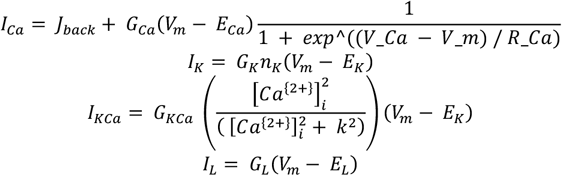

The Nernst potential for calcium is given by:

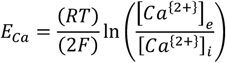

where R is the gas constant, T is the temperature, F is the Faraday constant, [*Ca*^{2+}^ ]_*e*_ and [*Ca*^{2+}^]_*i*_ are the extracellular and intracellular calcium concentrations.

### B. Mathematical Derivation of the Bidomain Model

To model the effects of the potential difference across the cell membrane, the tissue is divided into two continuous domains: the intracellular and extracellular spaces. These domains together represent the complete volume of the uterine muscle, and they are electrically coupled through the cell membrane [4]. The intracellular domain is considered continuous because the muscle cells are connected by gap junctions, allowing for electrical signal propagation [6].

The transmembrane potential *V*_*m*_ is defined as the difference between the intracellular potential *u*_*i*_ and extracellular potential *u*_*e*_:

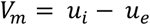

This potential difference drives ionic currents across the cell membrane, generating action potentials. The current densities in the intracellular and extracellular domains are:

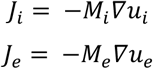

where *J*_*i*_ and *J*_*e*_ are the intracellular and extracellular current densities, and *M*_*i*_ and *M*_*e*_ are the conductivity tensors for the intracellular and extracellular spaces, respectively.

The bidomain model views the uterus as two overlapping Ohmic conductors separated by the cell membrane, which is modeled as a capacitor. By applying the principle of current conservation and accounting for the capacitive nature of the membrane, the governing equations are:

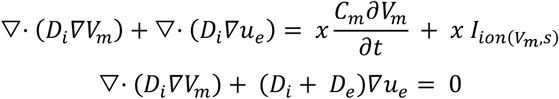

where *D*_*i*_ and *D*_*e*_ are the intracellular and extracellular conductivity tensors, *C*_*m*_ is the membrane capacitance, and 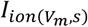 represents the ionic current. These equations describe the conservation of electrical current in the intracellular and extracellular spaces, accounting for the capacitive properties of the membrane and the ionic currents generated by cellular activity.

Boundary conditions impose no flux at the boundaries of the domain, ensuring that no current leaves the tissue:

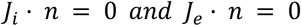

where *n* is the normal vector to the boundary. The primary unknowns in this system are the transmembrane potential *V*_*m*_ and the extracellular potential *u*_*e*_, while the cellular state variables s are governed by the action potential model in Section A.

### C. Numerical Solution of the Bidomain Equations

Where To numerically solve the bidomain equations, we implemented a second-order Strang splitting scheme, which separates the PDE system from the ODE system [12]. At each time step, the PDE system is discretized using linear finite elements defined on a 3D tetrahedral mesh of the uterine domain. The resulting algebraic system of equations is solved iteratively using a Krylov subspace method, which efficiently handles large, sparse matrices.

The finite element equations were derived using the Galerkin approach, which ensures stability and accuracy in the numerical solution. The forward solver was implemented using the *cbcbeat* library [13], built on the FEniCS [9, 16], Gotran [17], and dolfin-adjoint [18] frameworks. These tools allow for efficient handling of time-dependent PDEs and provide automated differentiation for adjoint-based optimization methods.

The Krylov solvers from PETSc [19] were employed to solve the large linear systems, ensuring fast and efficient computations. This approach allows us to handle the complex 3D uterine geometry and accurately model the propagation of electrical signals across the tissue.

### D. 3D Realistic Uterine Muscle Mesh

We used a 3D realistic mesh of the uterus provided by the FEMONUM project [10], derived from MRI images at 34.5 weeks of amenorrhea. The mesh was transformed into a volumetric representation by generating a tetrahedral mesh using TetGen [19]. This volumetric mesh captures the full thickness and structure of the uterine muscle, providing a realistic framework for simulating electrical signal propagation.

To account for regional variations in conductivity, we used a region-growing algorithm based on Poisson disk sampling [11]. This method ensures that seed points are well distributed across the mesh, preventing clustering of regions. The algorithm assigns a label to each region, which is propagated to neighboring vertices, ensuring that the entire mesh is segmented into distinct regions.

This segmentation allows for simulating anisotropic propagation of electrical signals, which is essential for accurately modeling the complex electrical behavior of the uterus during contractions.

## III. RESULTS

In this section, we present the results of the numerical simulations using the 3D volumetric mesh of the uterus. We run coupled cell model bidomain simulations with the Red3 electrophysiological model, as detailed earlier. The results capture the dynamic propagation of electrical signals throughout the uterine tissue, with specific focus on depolarization, repolarization, and the impact of isolated regions with different conductivities.

### A. Propagation of Electrical Activity

The simulation results over a 6-second interval reveal the spread of electrical activity initiated at the fundus of the uterus. In Figure 1, we observe the evolution of the electrical signal as it propagates through the uterine tissue. The series of panels (a) to (f) correspond to different time points in the simulation—specifically 1s, 2s, 3s, 4s, 5s, and 6s, respectively. The red regions indicate areas of depolarization, where the transmembrane potential rises, while the blue areas represent regions at resting potential. Figure 2 demonstrates the clear wave-like propagation of the depolarization wave starting from the fundus, spreading uniformly toward the lower regions of the uterus. This propagation pattern is critical for initiating coordinated contractions during labor.

**Figure 1:**
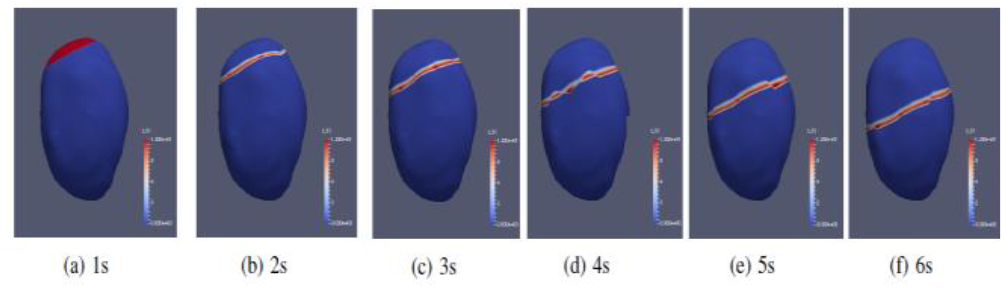
Results of the simulation panels (a) to (f) correspond to the electrical activity of the uterus. Each figure corresponds to a given time of the simulation: respectively 1s, 2s, 3s, 4s, 5s, 6s.

**Figure 2:**
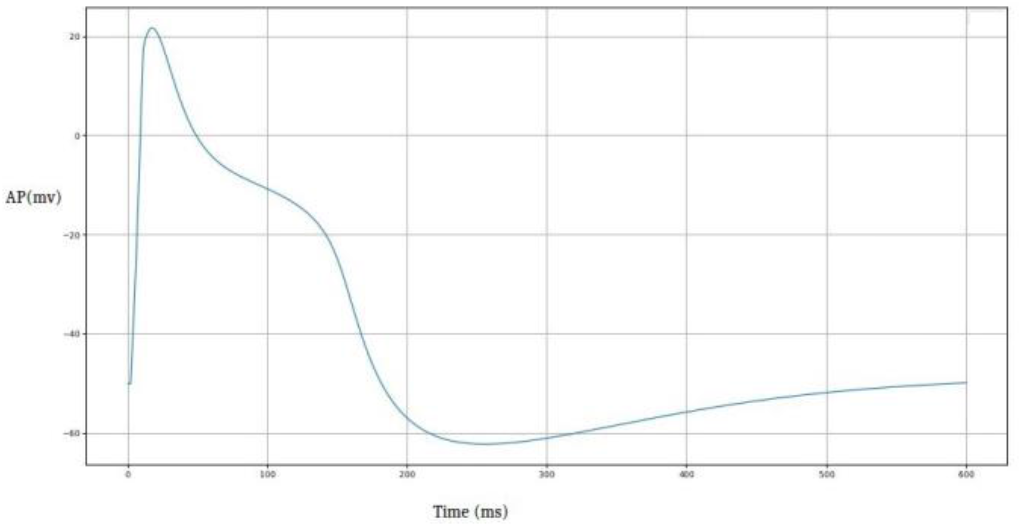
Action potential of a single uterine cell simulated with the Red3 model.

**Figure 3:**
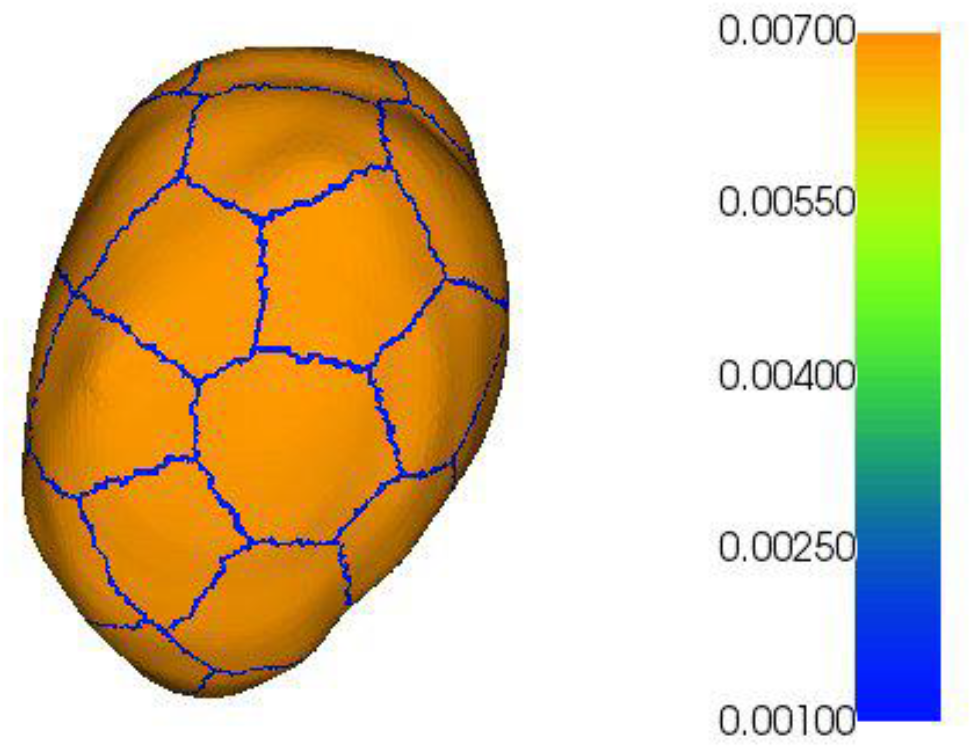
Illustration of 27 regions where the conductivities at the borders are 0.001 mS/cm and 0.0068 mS/cm inside the regions.

### B. Action Potential of a Single Cell

To further examine the electrical behavior at the cellular level, we simulated the action potential for a single uterine cell. The action potential represents the voltage changes across the cell membrane in response to ionic fluxes. Figure 2 shows the typical waveform of the action potential, where the initial depolarization phase leads to a rapid rise in potential, followed by a plateau, and then repolarization. The cell starts from a negative resting potential (around -50mV), rapidly depolarizes to a positive peak (above 20mV), and then gradually returns to its negative resting state. This dynamic process underlies the ability of uterine cells to generate and propagate electrical signals required for muscle contraction.

### C. Isolated Regions with Varied Conductivity

To simulate the effect of isolated regions in the uterine tissue, we divided the uterus into 27 distinct regions with different conductivity values. The conductivity inside the regions was set to 0.0068 mS/cm, while the conductivity at the borders was reduced to 0.001 mS/cm to simulate electrical isolation. Figure 4 shows a visual representation of the uterus segmented into these 27 regions, with a color map indicating the variation in conductivity. This segmentation helps to mimic the real-world heterogeneity of the uterus, where different regions may exhibit varying electrical properties. The simulation results demonstrate that regions with lower conductivity at the borders experience delayed signal propagation, highlighting the importance of regional electrical isolation in the overall coordination of uterine contractions.

## IV. DISCUSSION AND CONCLUSION

In this study, we presented a comprehensive mathematical framework for simulating the electrical activity of the human uterus using a bidomain model. Our approach integrates the Red3 electrophysiological cell model specifically designed to represent the ionic behavior in uterine smooth muscle cells, offering a detailed simulation of the propagation of electrical excitation throughout the uterus. By employing a non-linear reaction-diffusion type model, the simulation effectively captures the complex interactions governing uterine contractions during pregnancy and labor.

One of the key contributions of this work is the adoption of a continuous bidomain model, where the uterus is treated as two coupled domains—intracellular and extracellular spaces. This approach moves away from a discrete, cell-level perspective and instead focuses on the averaged potentials and currents at each point in space, which provides a more scalable and accurate representation of uterine tissue. The use of a conductivity tensor at each point allows for the preservation of the tissue’s heterogeneous structure, accounting for the anisotropy present in biological tissues. This is a significant improvement over previous models, which often relied on simpler, isotropic assumptions or geometric simplifications [1][2].

### A. Simulation Results and Their Implications

The numerical simulations performed on a three-dimensional volumetric mesh provide key insights into the propagation of electrical signals across uterine tissue. Our results demonstrate the ability of the developed code to accurately simulate both the excitation and repolarization phases of uterine electrical activity. The isotropic propagation of depolarization waves through the tissue was clearly captured, with the results aligning well with known physiological data on electrical signal propagation in the uterus [3].

The simulations revealed several important features of uterine electrical behavior. First, the depolarization wave initiates at the fundus and propagates downward in a coordinated manner, mimicking the natural excitation process observed during labor. Second, the repolarization phase, which follows the depolarization, restores the membrane potential to its resting state, ensuring the cells are prepared for subsequent contractions. The results further suggest that this method can be applied to simulate various pathological phenomena, such as asynchronous contractions or regions of electrical isolation, which may contribute to conditions like premature birth [4][5].

### B. Advantages of the Bidomain Approach

By employing the bidomain model, we were able to simulate the interaction between the intracellular and extracellular spaces, providing a more accurate representation of how electrical signals propagate through the uterus. This approach allows for a continuous exchange of ionic currents across the cell membrane, driven by transmembrane potential differences. The Red3 model’s inclusion of key ionic currents (e.g., calcium, potassium, and leakage currents) ensures that the simulated action potentials accurately reflect the underlying physiology of uterine smooth muscle cells [6].

Additionally, the bidomain approach offers several advantages over traditional single-domain models. First, it allows for the differentiation of intracellular and extracellular potentials, which is critical for capturing the complex coupling between these domains. Second, the use of a 3D volumetric mesh enables simulations to account for the full anatomical geometry of the uterus, providing insights into how electrical activity propagates across different regions of the organ. Finally, this method is flexible enough to be adapted for other types of excitable tissues beyond the uterus, making it a valuable tool for a wide range of biomedical applications [7][8].

### C. Applications to Pathological Uterine Phenomena

The developed simulation framework provides a solid foundation for investigating pathological conditions associated with abnormal uterine contractions. By adjusting the parameters of the model, such as the conductivities and ionic channel properties, it is possible to simulate scenarios that might lead to premature labor or uterine inertia. The ability to model isolated regions with different conductivities, as demonstrated in our simulations, allows for the investigation of how electrical isolation can disrupt normal contraction patterns and contribute to abnormal labor outcomes [9].

Furthermore, the model has the potential to be extended for use in predicting electrohysterograms (EHGs), which are non-invasive recordings of uterine electrical activity. By solving the forward problem in the physical domain of the uterus, the model can be used to simulate the generation of EHG signals, providing a valuable tool for predicting uterine behavior in both healthy and pathological states [10][11].

### D. Conclusion

In conclusion, the present study provides a robust framework for simulating the electrical activity of the human uterus using a bidomain approach. The developed code successfully calculates intracellular and extracellular potentials within a 3D volumetric mesh, accurately capturing the dynamics of action potentials and their propagation across the tissue. This method not only advances our understanding of normal uterine function but also provides a platform for investigating pathological phenomena that contribute to complications such as preterm birth.

The flexibility of the bidomain model, combined with the Red3 cellular model, allows for a wide range of applications in both clinical and research settings. Future work will focus on refining the model to include more detailed anatomical features and extending its application to personalized medicine, where patient-specific models of uterine electrical activity could be used to predict labor patterns and guide medical interventions

